# Comparative host-pathogen dynamics of Snake Fungal Disease in sympatric species of water snakes (*Nerodia*)

**DOI:** 10.1101/2022.03.08.483470

**Authors:** Stephen F. Harding, C. Guilherme Becker, Jessica Yates, Paul Crump, Michael R. J. Forstner, Stephen J. Mullin, David Rodriguez

**Affiliations:** Department of Biology, Texas State University, 601 University Drive, San Marcos, TX 78666, USA; Department of Biology, The Pennsylvania State University, 606 Mueller Laboratory, University Park, PA 16802, USA; Department of Biology, Stephen F. Austin State University, 2102 Alumni Drive, Nacogdoches, TX, 75962, USA; Texas Parks and Wildlife Department, Austin, TX, 78744, USA; Department of Biological Sciences, Arkansas State University, PO Box 599, State University, AR, 72467, USA

**Keywords:** Disease dynamics, *Ophidiomyces ophiodiicola*, Ophidiomycosis, Riparian habitat, Threatened Species

## Abstract

The ascomycete fungus *Ophidiomyces ophiodiicola* (*Oo*) is the causative agent of ophidiomycosis (Snake Fungal Disease), which has been detected globally. However, surveillance efforts in the central U.S., specifically Texas, have been minimal. The threatened and rare Brazos water snake (*Nerodia harteri harteri*) is one of the most range restricted snakes in the U.S. and is sympatric with two wide-ranging congeners, *N. erythrogaster transversa* and *N. rhombifer*, in north central Texas; thus, providing an opportunity to test comparative host-pathogen dynamics in this system. To accomplish this, we surveyed a portion of the Brazos river drainage (~400 river km) over 29 months and tested 150 *Nerodia* spp. for the presence of *Oo* via quantitative PCR and recorded any potential signs of *Oo* infection. We found *Oo* was distributed across the entire range of *N. h. harteri*, *Oo* prevalence was 46 % overall, and there was a significant association between *Oo* occurrence and signs of infection in our sample. Models indicated adults had a higher probability of *Oo* infection than juveniles and subadults, and adult *N. h. harteri* had a higher probability of infection than adult *N. rhombifer* but not higher than adult *N. e. transversa*. High *Oo* prevalence estimates (94.4%) in adult *N. h. harteri* has implications for their conservation and management owing to their patchy distribution, comparatively low genetic diversity, and threats from anthropogenic habitat modification.

## Introduction

Emerging infectious diseases (EIDs) have become a serious threat to wildlife with several outbreaks across broad taxonomic groups ^1^. Prime examples of fungal infections in wildlife across the globe include chytridiomycosis caused by *Batrachochytrium dendrobatidis* ^2–4^ or *B. salamandrivorans* ^5–7^ in amphibians, as well as white-nose syndrome caused by *Pseudogymnoascus destructans* infections in bats ^8–11^. Similarly, the ascomycete fungus *Ophidiomyces ophiodiicola* (*Oo*) is the causative agent of ophidiomycosis (Snake Fungal Disease), an EID in North America and Europe that was also recently detected in Asia ^12–15^. Signs of infection first appear as dermatitis, and if not cleared, infections will advance to lesions, ulcers, and tissue necrosis—the latter of which can precede death ^12,16^.

In North America, *Oo* was first detected in 2008 among wild populations of eastern massasaugas (*Sistrurus catenatus*) from Illinois ^14^. At the population level, in combination with low genetic diversity and climatic factors, *Oo* infections were later implicated in the decline of an isolated population of timber rattlesnakes (*Crotalus horridus*) in New Hampshire ^12,17^. The pathogen has since been detected across the contiguous United States in numerous species throughout the Midwest, East Coast,^18–22^ California, ^23^ Arizona (E. Nowak, unpublished data), and most recently, Texas. Prior to this study, two reports from the Texas Parks and Wildlife Department confirmed *Oo* infections in *Nerodia harteri harteri*, the Brazos water snake (or Brazos River Watersnake), in 2016 and in an Eastern Patch-nosed Snake (*Salvadora grahamiae*) in 2017 (Dryad repository: https://doi.org/XX.XXXX/dryad.XXXXXXX). Owing to a lack of surveillance efforts, however, little is known regarding the host-pathogen dynamics of ophidiomycosis in the state.

Confirmed infections of *Oo* across multiple snake genera suggest low host specificity 22,24. Thus, this pathogen presents a potential concern regarding biodiversity and conservation in Texas, as the state harbors the highest diversity of snakes in the U.S. ^25–28^ with 76 documented species and 115 varieties—if subspecies are included—eight of which are considered threatened 29. *Nerodia harteri* is among those species and also is one of two snakes endemic to Texas. Both of its subspecies *N. h. harteri and N. h. paucimaculata*, the Concho water snake (or Concho River Watersnake)^30^, show high affinity for water bodies and only inhabit disjunct stretches of the Brazos and Colorado River basins, respectively ^31–33^.

Historically, both subspecies of *N. harteri* have been affected by anthropogenic influences such as habitat destruction and variable water flow regimes from dam releases ^32,34^. *Nerodia h. paucimaculata* was listed as federally threatened in 1986 ^35^ and then delisted in 2011 owing to a recovery in their populations ^36^. *Nerodia h. harteri* was initially petitioned for federal listing in 1984 ^36^ and, although the U.S. Fish and Wildlife Service found that listing was not warranted after a 12-month review in 1985 ^37^, the snake remained on the candidate species list until 1994 ^38^. More recent studies detected declines in overall abundance and evidence of low genetic diversity in *N. harteri* ^31,34^. In contrast, two non-threatened sympatric congeners, Diamond-backed water snakes (*N. rhombifer*) and Blotched water snakes (*N. erythrogaster transversa*), have much larger ranges ^30,39^, attain larger sizes, are more fecund, and are more common ^40^; thus, providing a suitable system to test differences in host-pathogen dynamics among species that differ in their ecological attributes and conservation status.

Factors that contribute to the probability of *Oo* infection and mortality among individuals and species vary considerably, although some studies have shown climate and season are associated with incidence and severity of infection ^12,17,18,41–43^. However, interspecific ^44^ and demographic factors ^18^ associated with pathogen prevalence are still understudied. One study presented evidence of interspecific *Oo* prevalence differences in Georgia (USA) ^44^, and another documented vertical transmission of *Oo* from infected mothers to neonates among viviparous and oviparous snakes ^45^. However, there are no published studies assessing the role of interspecific and demographic factors in the distribution of *Oo* in Texas snake populations. Assessments of *Oo* prevalence (i.e., the proportion of *Oo-*positive snakes among a subset of a population), geographic distribution of the pathogen, and pathogen-associated abiotic (i.e., temperature and rainfall) and biotic factors (i.e., species and demography) are needed in this region—especially among endemic species with small populations like *N. harteri* ^17^.

We address the paucity of surveillance data for *Oo* in Texas by conducting the first population-level survey of *Oo* prevalence among *Nerodia* species within the Brazos River basin. Our goals were to: 1) specifically survey areas with historically abundant populations of *N. h. harteri*; 2) estimate the geographic distribution of *Oo* in the upper Brazos river basin; and 3) test for associations between biotic and abiotic factors and *Oo* infection prevalence estimates among *Nerodia* inhabiting those areas. We predicted that *N. h. harteri* would be more susceptible to *Oo* infections on account of their comparatively smaller population sizes. Our study seeks to increase understanding of the distribution of *Oo* in this region of the U.S. and comparative *Oo* infection dynamics among sympatric *Nerodia*. We also aim to provide key ecological and epidemiological data for the future management of threatened *N. h. harteri* populations.

## Results

In total, we captured, visually inspected, and swabbed 150 snakes (Table 1). We collected 76 *N. rhombifer* (33 males, 41 females, 2 undetermined sex), 34 *N. e. transversa* (11 males, 15 females, 8 undetermined sex), and 40 *N. h. harteri* (17 males, 20 females, 3 undetermined sex). During our surveys, we recaptured six snakes: one *N. rhombifer*, four *N. e. transversa*, and one *N. h. harteri*. The *N. rhombifer* and *N. e. transversa* were sampled, released, and recaptured during the same sampling trip, and the single *N. h. harteri* was recaptured 14 days after the initial capture. The average snout-vent length (SVL) for *N. rhombifer*, *N. e. transversa*, and *N. h. harteri* was 63.78 cm (SD = 30.06), 51.20 cm (SD = 21.42), and 40.31 cm (SD = 17.16), respectively. The average mass for *N. rhombifer*, *N. e. transversa*, and *N. h. harteri* was 249.26 g (SD = 231.20), and 143.78 g (SD = 134.17), and 62.30 g (SD = 62.22), respectively.

**Table 1.**
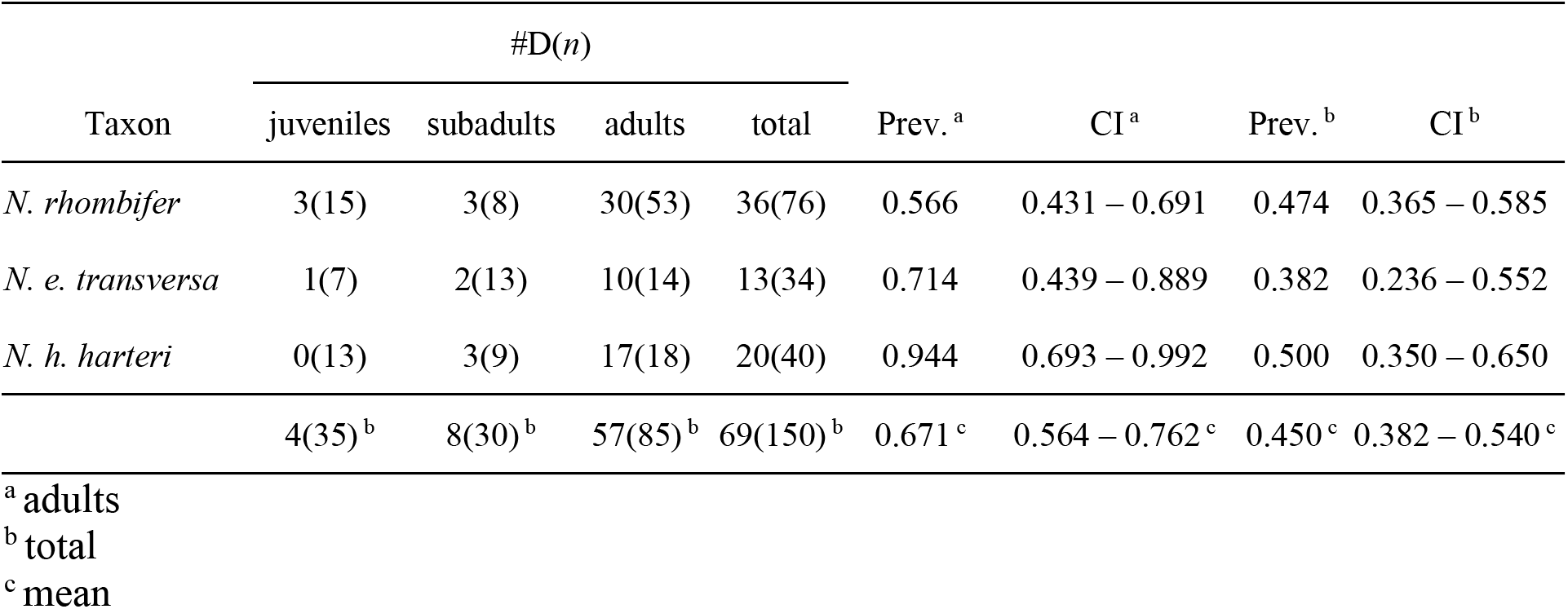
Number of *Nerodia* positive for *Ophidiomyces ophiodiicola* (#D) among the total collected (*n*), the estimated prevalence as a proportion for the total number of each species collected (Prev.), and the 95% binomial confidence intervals for each prevalence estimate (CI).

| Taxon | #D(*n*) |  |  |  | Prev. <sup>a</sup> | CI <sup>a</sup> | Prev. <sup>b</sup> | CI <sup>b</sup> |
| --- | --- | --- | --- | --- | --- | --- | --- | --- |
|  | juveniles | subadults | adults | total |  |  |  |  |
| *N. rhombifer* | 3(15) | 3(8) | 30(53) | 36(76) | 0.566 | 0.431 – 0.691 | 0.474 | 0.365 – 0.585 |
| *N. e. transversa* | 1(7) | 2(13) | 10(14) | 13(34) | 0.714 | 0.439 – 0.889 | 0.382 | 0.236 – 0.552 |
| *N. h. harteri* | 0(13) | 3(9) | 17(18) | 20(40) | 0.944 | 0.693 – 0.992 | 0.500 | 0.350 – 0.650 |
|  | 4(35) <sup>b</sup> | 8(30) <sup>b</sup> | 57(85) <sup>b</sup> | 69(150) <sup>b</sup> | 0.671 <sup>c</sup> | 0.564 – 0.762 <sup>c</sup> | 0.450 <sup>c</sup> | 0.382 – 0.540 <sup>c</sup> |

<sup>a</sup> adults
<sup>b</sup> total
<sup>c</sup> mean

We detected potential signs of infection (SOI, see Methods for definition; Fig. 1) on 30 *N. rhombifer* (26 adults, 1 subadult, 3 juveniles), 8 *N. e. transversa* (5 adults, 3 subadults), and 13 *N. h. harteri* (11 adults, 2 subadults). We detected *Oo* on 39 of the 51 snakes with SOI (20 *N. rhombifer*, 6 *N. e. transversa*, and 13 *N. h. harteri*). We also detected *Oo* on 30 snakes not exhibiting any apparent SOI (16 *N. rhombifer*, 7 *N. e. transversa*, and 7 *N. h. harteri*) (Table 1). We detected *Oo* on 11 of the 13 snakes with SOI that were subset for skin biopsies. When we compared *Oo* detections from swab and biopsy extractions, there were 11 out of 11 detections from swab extractions and 9 out of 11 detections from the skin biopsy extractions. Both swab and skin biopsy qPCR tests agreed on the two negatives that were observed in this subset.

**Figure 1.**
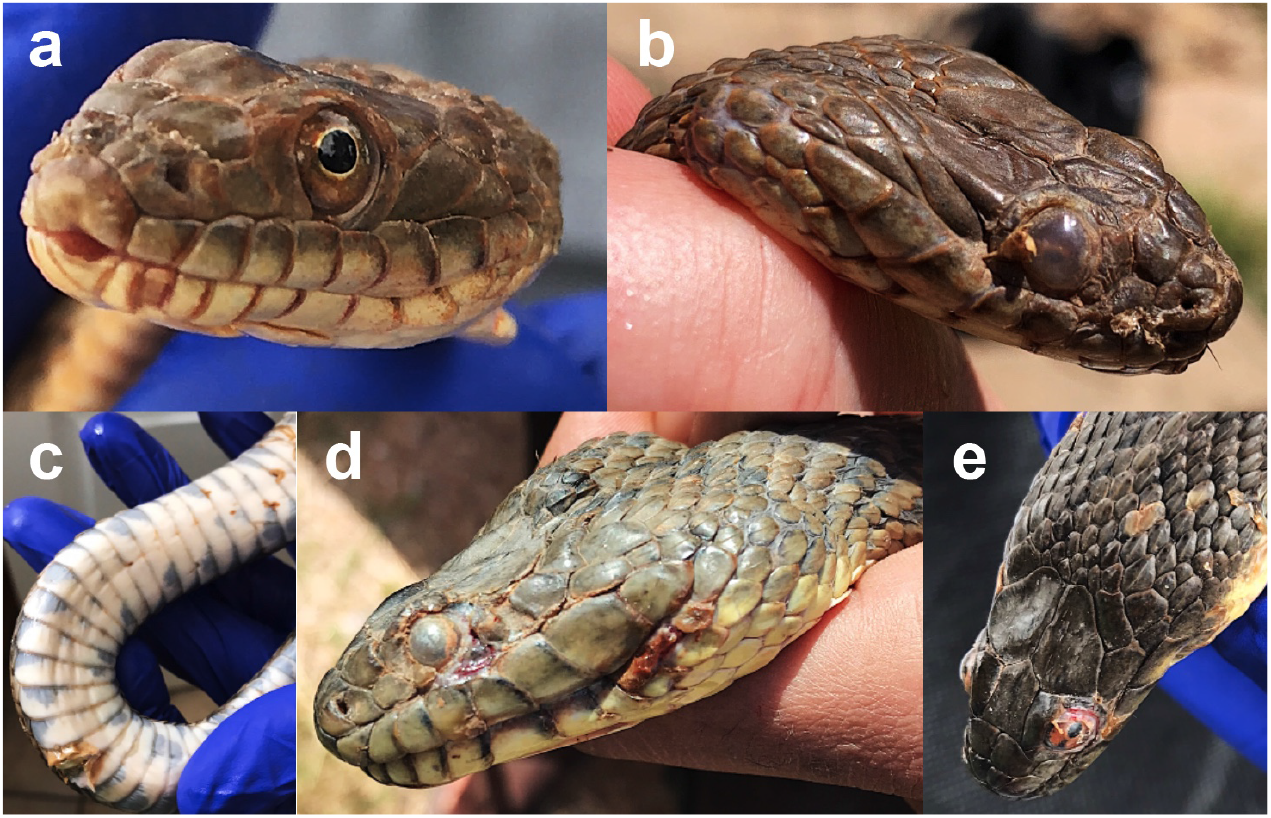
Varying signs of *Ophidiomyces ophiodiicola* infections. Lesions in *Nerodia harteri harteri* (panels a and b) and *N. rhombifer* (panels c, d, and e) were observed during a 29-month surveillance effort in the upper Brazos River basin (Texas, U.S.A.).

Our estimated *Oo* prevalence for *N. rhombifer*, *N. e. transversa*, and *N. h. harteri* was 47.4 % (CI: 36.5 – 58.5 %), 38.2 % (CI: 23.6 – 55.2 %), and 50.0 % (CI: 35.0 – 65.0 %), respectively, and overall, our uncorrected *Oo* prevalence was 46.0% (CI: 38.2 – 54.0 %) (Table 1). The false-negative rate was 0 for swab samples and 0.153 for the skin biopsies, and the Bayesian approximation of corrected *Oo* prevalence was 48.2 % (CI: 40.5 – 56.4 %) among sampled *Nerodia* species. Adult *N. h. harteri* showed the highest estimate of *Oo* prevalence (94.4 %; CI: 69.3 – 99.2 %; Fig. 2; Table 1). We captured more snakes in May and June during their peak activity (Fig. 3).

**Figure 2.**
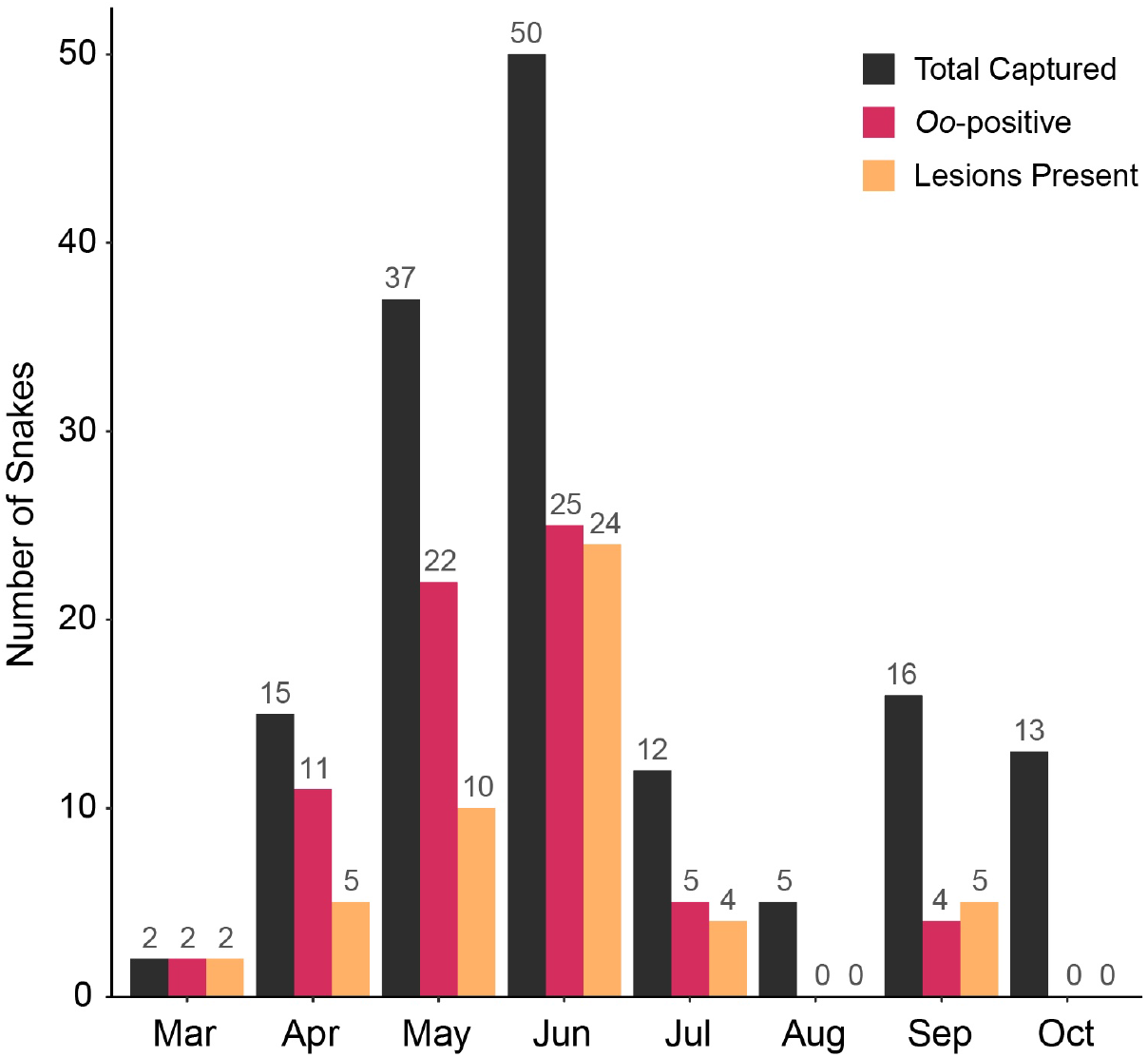
Number of wild-caught adults and total sample size for three *Nerodia* species tested for the presence of *Ophidiomyces ophiodiicola* via qPCR indicated by gray bars. Closed circles represent estimates of prevalence for each species and a subset of adults, with black lines showing the 95% confidence intervals.

**Figure 3.**
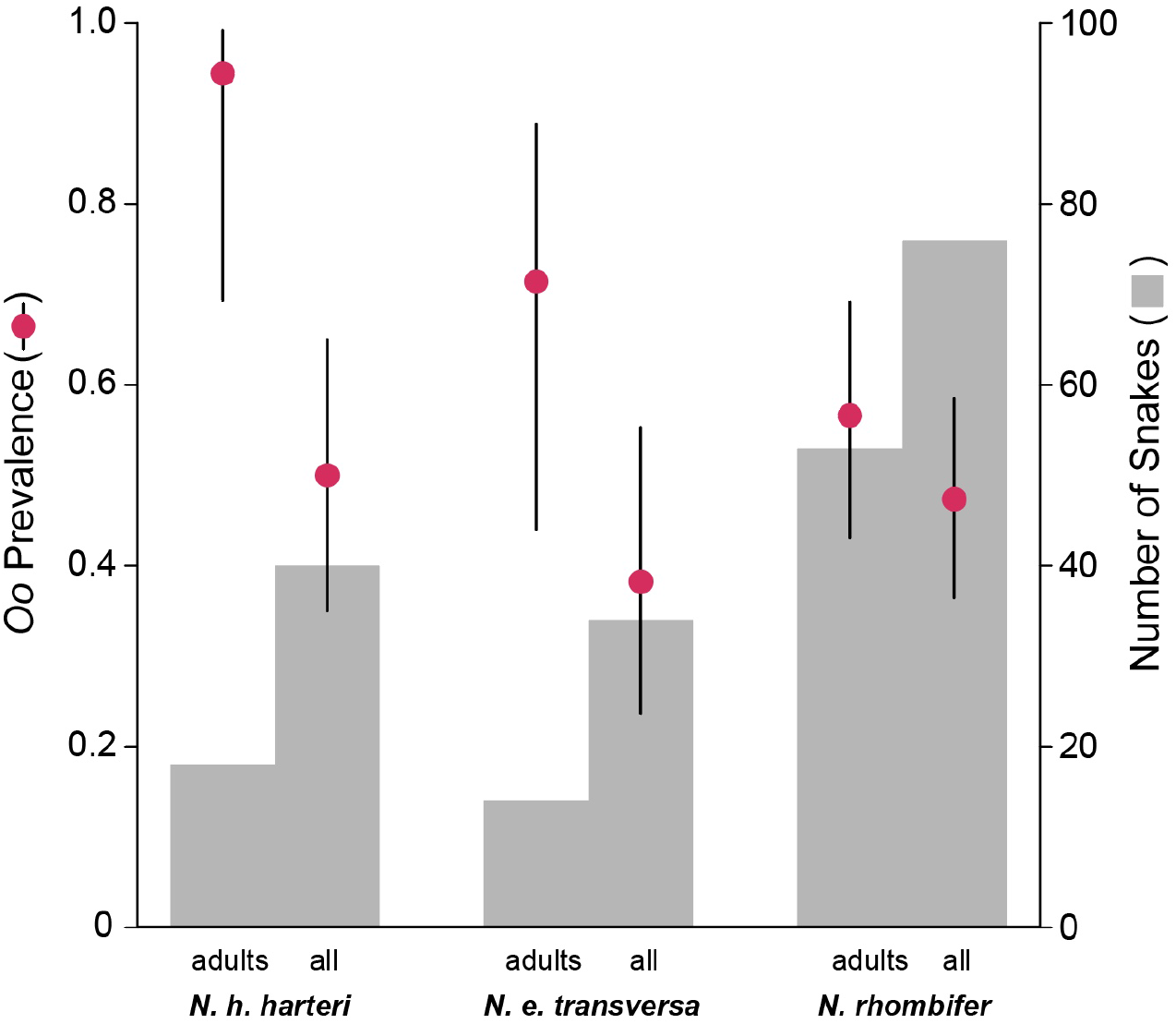
Temporal histogram of the total number of snakes captured, the number of snakes that tested positive for *Ophidiomyces ophiodiicola* via qPCR, and the number of snakes showing signs of infection. Surveillance efforts were performed from 16 April 2018 through 6 October 2020.

Geographically, we detected *Oo* at 13 of the 21 sites (Fig. 4) and in all eight counties sampled. Our county-level prevalence estimates ranged from 25.0 % (Hill Co.) to 75.0 % (Bosque Co.) (Table 2). We found a significant association between signs of infection and *Oo* occurrence (Fishers Exact Test, *P* < 0.001) but not between species and SOI (*P* > 0.20) nor between sex and *Oo* occurrence (*P* > 0.40). We also detected significant differences in body condition (BCI) values among adult *N. e. transversa* with and without signs of infection (Kruskal-Wallis test, *P* < 0.01; Table 3).

**Table 2.** Texas counties sampled for *Nerodia rhombifer*, *N. e. transversa*, and *N. h. harteri*, the number of snakes that tested positive for *Ophidiomyces ophiodiicola* (#D) among the total collected in each county (*n*), the estimated prevalence as a proportion (Prev.), and the 95% binomial confidence intervals for each estimate (CI).

| County | #D(*n*) | Prev. | CI |
| --- | --- | --- | --- |
| Bosque | 3(4) | 0.750 | 0.238 – 0.966 |
| Haskell | 8(12) | 0.667 | 0.376 – 0.870 |
| Hill | 2(8) | 0.250 | 0.063 – 0.623 |
| Hood | 24(46) | 0.522 | 0.379 – 0.660 |
| Jones | 3(6) | 0.500 | 0.168 – 0.832 |
| Palo Pinto | 14(33) | 0.424 | 0.270 - 0.595 |
| Shackelford | 12(34) | 0.353 | 0.213 - 0.524 |
| Stephens | 3(7) | 0.429 | 0.144 - 0.770 |
| Overall | 69(150)<sup>a</sup> | 0.460<sup>b</sup> | 0.382 – 0.540<sup>b</sup> |

<sup>a</sup> total
<sup>a</sup> mean

**Table 3.** Summary statistics from Kruskal-Wallis One Way ANOVA tests (TS = test statistic; df = degrees of freedom) for differences in body condition (BCI) when considering *O. ophiodiicola* occurrence (*Oo-occ*) and signs of infection (SOI) across life stages of *Nerodia rhombifer*, *N. e. transversa*, and *N. h. harteri*. Significant results are in bold type.

|  | *N. h. harteri* |  |  | *N. e. transversa* |  |  | *N. rhombifer* |  |  |
| --- | --- | --- | --- | --- | --- | --- | --- | --- | --- |
|  | TS | df | *P*-value | TS | df | *P*-value | TS | df | *P*-value |
| Adult BCI x *Oo-occ* | 0.232 | 1 | 0.630 | 1.029 | 1 | 0.311 | 0.295 | 1 | 0.587 |
| Adult BCI x SOI | 0.284 | 1 | 0.594 | **6.942** | **1** | **0.008** | 2.995 | 1 | 0.084 |
| Sub-adult BCI x *Oo-occ* | 0.600 | 1 | 0.439 | 2.262 | 1 | 0.132 | 0.125 | 1 | 0.724 |
| Sub-adult BCI x SOI | 0.343 | 1 | 0.558 | 0.077 | 1 | 0.782 | 1.000 | 1 | 0.317 |
| Juvenile BCI x *Oo-occ* | - | - | - | 0.250 | 1 | 0.617 | 0.521 | 1 | 0.471 |
| Juvenile BCI x SOI | - | - | - | - | - | - | 0.750 | 1 | 0.387 |

**Figure 4.**
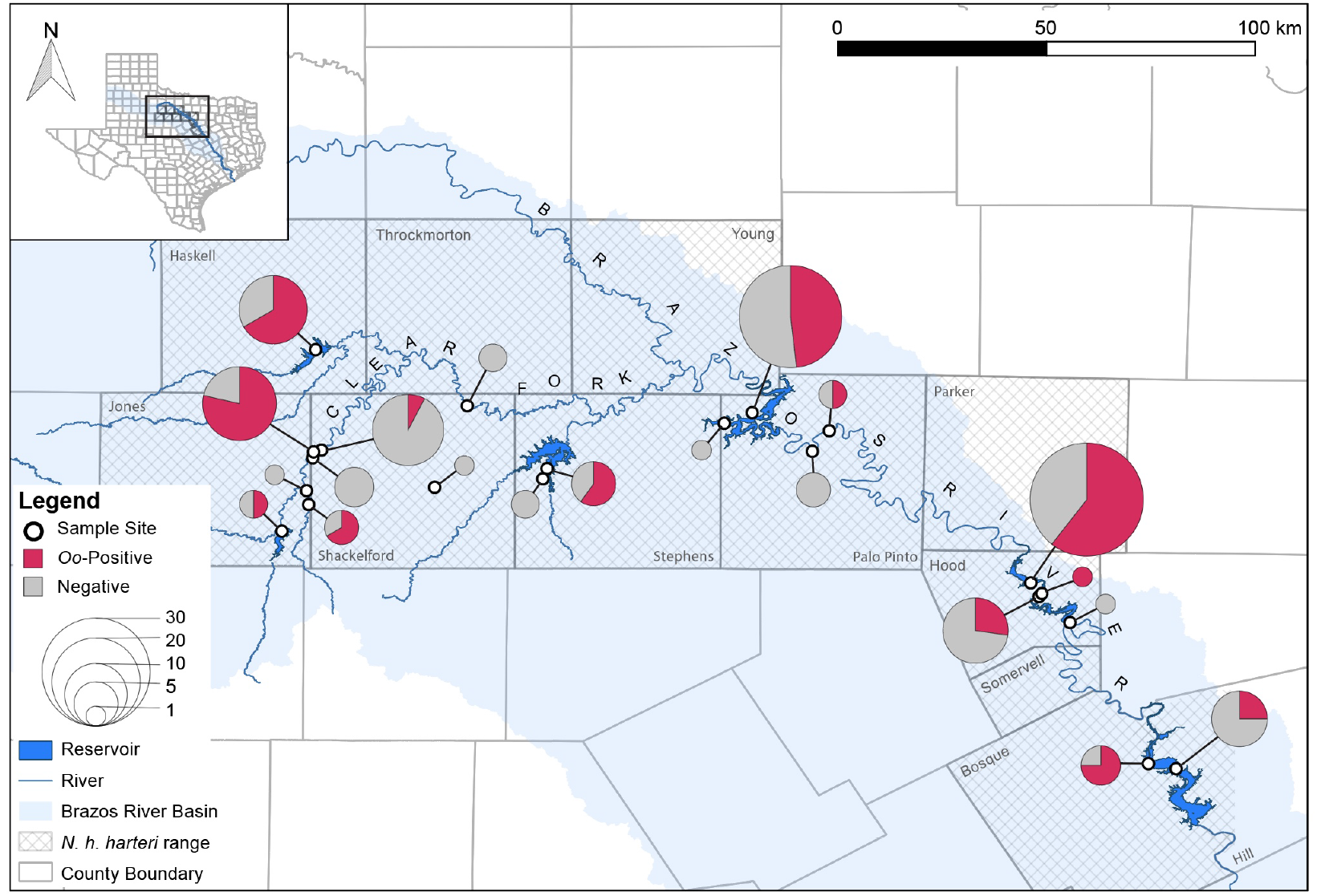
Map of sampling sites demarcated with open circles. Cross-hatching indicates the historical range of *Nerodia harteri harteri* at the county level within the Brazos River drainage designated by the shaded area. Each pie chart represents the relative proportion of *Nerodia* that tested positive or negative for *Ophidiomyces ophiodiicola* via qPCR. The size of the pie chart represents the relative number of snakes collected per site.

The most parsimonious model explaining *Oo* occurrence included species and life stage with temporal autocorrelation accounted for by sample month as an AR1 random effect (Supplementary Table S1). The model indicated that *N. h. harteri* were more likely to be infected than *N. rhombifer* (β = –1.28; SE = 0.64), while juvenile snakes (β = −2.87; SE = 0.66) and subadults (β = −2.00; SE = 0.57) were less likely to be infected than adults (Supplementary Table S2; Supplementary Table S3).

Our modeled average probability of infection for *N. rhombifer*, *N. e. transversa*, and *N. h. harteri* was 20.7 % (CI: 9.20 – 40.3 %), 25.7 % (CI: 11.1 – 48.9 %), and 48.4 % (CI: 25.9 – 71.6 %), respectively (Supplementary Table S4). The modeled average probability of infection at the juvenile, subadult, and adult stage was 11.2 % (CI: 4.0 – 27.7 %), 23.2 % (CI: 9.6 – 46.2 %), and 69.1 % (CI: 48.9 – 83.9 %), respectively (Supplementary Table S5). Adult *N. h. harteri* had the highest value of estimated probability of infection (82.7%, CI: 60.9 – 93.6 %) (Supplementary Table S6).

## Discussion

Ophidiomycosis has been increasingly detected in North America across a broad range of host species of snakes, which includes some taxa of conservation concern ^14,17,24,46^. The initiation of pathogen surveillance efforts is an important step in determining the host-pathogen dynamics among potentially susceptible taxa. In this study, we addressed a major wildlife disease surveillance gap in Texas snakes by focusing on the upper Brazos River drainage, because it hosts the endemic and threatened Brazos water snake. To provide comparative *Oo* prevalence data, we also sampled two more common and widespread sympatric congeners. Owing to the consequences of small population sizes, we predicted that *N. h. harteri* could be more susceptible to *Oo* infections; thus, we used estimates of *Oo* prevalence and associated SOI among these sympatric species to test this prediction. Previous *Oo* surveys have reported prevalence estimates ranging from 1.8 to 66.5 % ^18,47–50^. We observed moderate prevalence across all samples (46.0 %) as well as at the species level (38.2 – 50.0 %). Prevalence estimates among adults were higher in *N. h. harteri* than in *N. rhombifer*, which partially supports our prediction regarding the greater susceptibility of *N. h. harteri* to *Oo* infections.

### Detection of ophidiomycosis

Signs of infection were a strong indicator of *Oo* occurrence, which confirms *Oo* infections in *Nerodia* populations from the upper Brazos River and its tributaries, and is potentially more widespread in this region given the range size of *N. rhombifer* and *N. e. transversa*. Even though we did not detect snakes that were moribund owing to severe infections in this population sample, some exhibited signs of an advancing infection (see Fig. 1). We have, however, also confirmed the presence of *Oo* on emaciated, moribund snakes from other parts of Texas—specifically, *Coluber constrictor*, *N. e. transversa*, and *Agkistrodon piscivorus* (Dryad repository: https://doi.org/XX.XXXX/dryad.XXXXXXX). Therefore, we are confident that the advanced infections observed in our study area likely represent progressing ophidiomycosis.

Subsampling for histopathological detection of *Oo* infections, and confirmation of ophidiomycosis, would be useful, albeit not tractable for *N. h. harteri* owing to its low abundance, conservation status, and the lack of detectable moribund individuals in this study. We have demonstrated that using a single swab with multiple strokes across the body is cost-efficient and suitable for distinguishing between SOI owing to *Oo* or other reasons, and serves as an alternative to using multiple swabs per snake. The use of multiple swabs decreases the probability of false-negatives ^48^, however, and would be beneficial in populations with lower *Oo* prevalence than we estimated in this region. In 12 snakes that exhibited potential SOI but did not test positive for the presence of *Oo*, it is likely those represent an injury, a cleared infection, other infection type, or a false negative. Considering our estimated false-negative rate for these data (~15 %), we can assume the number of true false-negatives in this sample is relatively low (2 samples), and the estimate of previously injured but not *Oo* infected snakes is approximately 6.70 % in our overall sample.

### Abiotic and biotic predictors of *Oo* detection

With the exception of adult *N. e. transversa*, our analyses did not detect differences in BCI between snakes with or without the presence of *Oo*, or with or without signs of infection (Table 3). These results are consistent with other reports that compared body condition among infected and non-infected snakes in laboratory settings ^16^ and in the wild ^18,47^. On the other hand, some studies have shown general body condition decreased with increased severity of ophidiomycosis ^14,41,43^. We did not observe that general pattern, which might be an artifact of the aquatic habitat these species use leading to decreased encounters of moribund or dead individuals. Additionally, capturing dying snakes is presumably difficult, as they are less active than healthy snakes. For instance, behavioral observations of wild eastern massasauga rattlesnakes showed that infected individuals moved less and remained concealed more often than their uninfected counterparts ^51^. Conversely, other studies of free-ranging pygmy rattlesnakes (*Sistrurus miliarus*) showed severity of *Oo* infection fluctuated and was not related to the probability of capture ^41^. Therefore, the lack of moribund individuals in our sample could result from disease induced changes in behavior or, alternatively, simply attributable to the stochastic nature of encounters in dynamic aquatic habitats.

Our results showed adult snakes were more likely to be infected than subadults and juveniles (Fig. 2; Supplementary Table S5). In general, *Nerodia* emerge from hibernation in early spring and are active through early summer ^39^; thus, we consider it possible that seasonal patterns and reproductive behavior might have contributed to age-class differences in *Oo* prevalence. We captured the majority of snakes—infected and non-infected—during the spring with captures decreasing during the summer, which is consistent with their behavior. We sampled snakes with lesions as early as March (Fig. 3), before the mating season. *Nerodia* mate in mid to late spring and parturition occurs during late summer to early fall ^39,52–54^. Therefore, the infected snakes we captured in March and April likely represent environmental transmissions ^12^, whereas infected snakes collected in June might have acquired infections via environmental transmission or contact with other infected snakes during reproduction events throughout April and May. The lower infection prevalence among the subadults and juveniles might have resulted from insufficient time in the environment to acquire infections and the lack of secondary contact during reproduction events. Although, a previous study has reported variable neonate mortality associated with *Oo* infection vertically transferred from the dam ^45^. Thus, it is also possible that infected neonates are being removed from the population before an *Oo* infection can be detected.

Even though we did not include season as a factor in our analyses, generally, our results are consistent with other surveys for *Oo* that suggest infected snakes are more likely to be encountered during the spring owing to fungal exposure during brumation through the winter months ^18,41,49,55^. These observations highlight concerns that climatic factors (i.e., cooler, wetter weather) could act synergistically with *Oo* and increase the severity of infection ^17,43^. Yet, mean monthly low temperatures and mean monthly precipitation were less parsimonious explanations to *Oo* occurrence in this study than species and life stage. The differences in *Oo* infection and severity among Texas snakes from different habitat types (i.e., terrestrial vs. aquatic) has yet to be tested; thus, expanded long-term surveillance of *Oo* infections and severity across years and habitat types will be informative in detecting patterns of seasonality.

### Host identity

Consistent with our observations, other studies have shown that *Oo* prevalence differs intra-generically among *Nerodia* ^44^. In our study, *N. h. harteri* exhibited higher *Oo* prevalence compared to *N. rhombifer*, which is concerning owing to the conservation status and rarity of *N. h. harteri*. With respect to *N. h. harteri* and *N. e. transversa*, the binomial 95% confidence intervals for *Oo* prevalence estimates overlapped, which could be interpreted as non-significance (Fig. 2); however, their relative abundance and distribution should be taken into consideration. For example, field surveys have reported that *N. h. harteri* were found in only ~300 km of stream and in two reservoirs within the Brazos River Drainage ^32,34^. These reports concluded that both subspecies of *N. harteri* are amongst the most range-restricted snakes in the U.S. In contrast, the range of *N. rhombifer* spans 12 states in the U.S. and 9 states in Mexico ^39,56^, and *N. e. transversa* is found in much of the eastern U.S. ^39^. Both snakes are generally more frequently encountered than *N. h. harteri*, which we captured at only 7 of 21 sites. This patchy distribution is congruent with estimates from previous studies ^31,32,34^. Comparatively, we captured *N. rhombifer* at 15 of 21 sites and *N. e. transversa* at 14 of 21 sites (Supplementary Fig. S1). Consequently, achieving captures at a rate needed to confer narrow confidence intervals for *Oo* prevalence estimates is difficult for relatively rare species like *N. h. harteri*. Yet, the number of *N. h. harteri* we captured during our survey (*N* = 40) was consistent with previous population surveys conducted over similar time intervals ^31,34^; therefore, uncertainty of *Oo* prevalence estimates could also be remedied by long-term surveillance efforts for this taxon.

### Conservation implications

Our survey data also suggest that *N. h. harteri* might now inhabit a more limited area within its historical range (Supplementary Fig. S1). Combined with a moderately high prevalence of *Oo* (47.4% overall, Table 1), this finding highlights the need for continued *Oo* surveillance and renewed conservation action planning. Previous studies have shown that both subspecies of *N. harteri* have low genetic diversity, and *N. h. paucimaculata* populations exhibit bottleneck signatures ^31,57^. The application of additional sampling and genetic markers are needed, however, to facilitate robust tests for population contractions and their timing in *N. harteri* ^31,57^. For instance, spatial ecology studies have shown *N. h. paucimaculata* can exhibit high site fidelity and are unlikely to move more than 1 km unless driven by stochastic factors such as variable water flow ^33^. It is likely that *N. h. harteri* movement among sites is likewise rare, given their low detection probability, the barriers to movement within the Brazos River (i.e., dams or unsuitable habitat), and evidence of genetic population structure in *N. h. paucimaculata*, which shares many ecological attributes with *N. h. harteri* ^31,57^.

Low genetic diversity, low dispersal potential, and high site fidelity are likely to increase inbreeding in populations of *N. harteri*. Inbreeding depression has been documented and implicated as a viable threat to snakes ^58^ and in other systems ^59–63^. Both inbreeding depression and an infection consistent with ophidiomycosis were attributed to population declines of timber rattlesnakes in New Hampshire ^17^. Estimates of mortality among snakes infected with *Oo* vary ^12,14,41,42^, and the contribution from inbreeding depression to increased risk of mortality from ophidiomycosis is still unknown. Conservation stakeholders should consider that inbreeding depression might act synergistically with other reported natural and anthropogenic stressors contributing to population declines in *N. h. harteri* and indeed, *N. harteri sensu lato*.

Our results have implications regarding the conservation status of *N. h. harteri* and indicate that ophidiomycosis is present in Brazos water snake populations. In the early 1980s, a proposal was submitted to the U.S. Fish and Wildlife Service to list both subspecies of *N. harteri* as endangered or threatened owing to their endemism and habitat loss ^64^. However, it was not listed because the snake was consistently present in suitable habitat within reservoirs ^35,37^. A population survey conducted from 2006–2008 reported the absence of snakes in areas where they were once historically abundant and concluded that interspecific competition, altered flow regimes, and negative effects from invasive species were likely contributing factors to low abundance ^34^. Currently, *N. h. harteri* is listed by the state of Texas as threatened ^29^, has a G1 (Critically) imperiled status (i.e., a very high risk of extinction ^65^), and a Near Threatened status from the IUCN ^66^. We have shown that ophidiomycosis is another potential threat to *N. h. harteri* populations and possibly other snake species in Texas—especially those with small population sizes. Thus, also initiating *Oo* surveillance in the Colorado River drainage among populations of *N. h. paucimaculata* is necessary to elucidate the host-pathogen dynamics for this species as a whole.

We marked snakes to avoid artificial inflation of snake counts and skewed estimates of pathogen prevalence, but this study was not designed to explicitly track disease outcomes. Thus, we cannot derive conclusions regarding neutral or negative population-level effects of ophidiomycosis among the few individuals we recaptured within short timeframes (<15 days).

We have, however, highlighted the need for long-term mark-recapture studies within infected populations to determine the impacts of ophidiomycosis in these and other Texas snake populations, such as a two-year study conducted for pygmy rattlesnakes in Florida ^41^. Texas encompasses broad ecological regions, each with unique vegetation cover and climatic patterns. Therefore, statewide surveillance will be useful in evaluating the wider distribution of the pathogen and assessing risks to the other seven snake species currently threatened in the state (e.g., *Cemophora coccinea copei*, *Cemophora coccinea lineri* [Texas endemic], *Coniophanes imperialis*, *Drymobius margaritiferus*, *Leptodeira septentrionalis*, *Pituophis ruthveni*, and *Tantilla cucullata*). Additionally, the deeper temporal dynamics of *Oo* in Texas are unclear. A retrospective survey of preserved specimens—similar to those conducted in eastern massasaugas 67 and other fungal pathogen systems ^68,69^—could also better elucidate seasonal and historical trends of *Oo* prevalence in Texas snakes, which remain largely understudied at the population-scale in this region of North America.

## Methods

### Study Area

In total, we sampled 21 sites distributed within Paint Creek, the Clear Fork of the Brazos, and sections of the Upper Brazos River basin (Fig. 4 and Supplementary Fig. S1); because of their patchy distribution and rarity, the majority of our effort focused on sites and counties where *N. h. harteri* populations were present historically ^31,32,34^. All procedures involving snakes were carried out in accordance with guidelines and regulations approved by the Texas State University Office of Research Integrity & Compliance (IACUC Protocol #26). Capture and sampling of snakes was approved by the Texas Parks and Wildlife Department (Scientific permits SPR-0316-059, SPR-0220-025, and SPR-0102-191), and the reporting in this manuscript follows the ARRIVE guidelines (https://arriveguidelines.org).

### Data Collection and *Oo* qPCRs

We collected snakes from 16 April 2018 through 6 October 2020. Our sampling trips were carried out opportunistically and depended on weather conditions without rainfall and with ambient air temperatures above 21°C; thus, we were unable to implement a structured temporal sampling design. Instead, we conducted the majority of our sampling trips during the spring and early summer months when *Nerodia* activity is known to be higher ^39,53^. Flooding in our study area also prevented surveys in June, July, and August of 2019, however, activity generally decreases after June. In 2020, surveys were delayed until May owing to mandatory travel restrictions attributable to the severe acute respiratory syndrome coronavirus 2 (SARS-CoV-2) outbreak.

To maximize the probability of capturing snakes present at a site, we utilized active and passive capture methods. Our active capture method consisted of kayaking navigable waterways and walking along shorelines to visually locate individuals and then capturing them by hand. Our passive capture method consisted of deploying Gee’s galvanized funnel crawfish traps (78.7 cm x 22.9 cm) along the shoreline ^33,70^ in areas identified as suitable *Nerodia* habitat (i.e., near chunk rock, edges of riffles, and large rip-rap). All traps were sanitized prior to deployment and after retrieval with a 1% bleach solution.

We handled each snake with fresh, sterile nitrile gloves. After capture, each snake was visually inspected for signs of *Oo* infection (see Fig. 1). We defined signs of *Oo* infection (SOI) as the presence of dermatitis, gross lesions, or any general scale abnormality (e.g., signs of inflammation, crust, nodules, or pus discharge) ^12,16^. We also photographed all potential visible signs of infection. To decrease costs, reduce handling time, minimize animal stress, and increase the surface area sampled, we swabbed the entire body of each snake using several strokes with a single sterile cotton-tipped swab (Medical Wire, MW113). We swabbed all surfaces of the head and then swabbed down the snake along the dorsum and the venter towards the tail. We immediately placed the swabs into a sterile 2 mL centrifuge tube with an O-ring screw cap. Of the snakes that exhibited SOI, we chose a subset (*N* =13) and collected skin biopsies (i.e., scale clips) from the affected area and placed the sample in 95% ethanol for downstream DNA extraction and qPCR testing. We kept tubes with swabs and skin biopsies on ice in the field and during transportation back to the laboratory, where they were stored at −20°C until processing.

After swabbing, we recorded sex, snout-vent length (SVL), and mass for each captured snake. To minimize cross contamination of equipment, we placed each snake in a sterile plastic bag prior to weighing it. We sterilized all non-disposable equipment with a 1% bleach solution prior to processing other individuals. We then marked the venter of each snake with a unique identifier using a cautery unit in case of future recapture ^71^. We used SVL to categorize individuals into species-specific life stages (see Supplementary Table S7) ^39,52–54,72,73^. For each county sampled, we retrieved monthly low temperature and monthly precipitation data from the National Centers for Environmental Information-National Oceanic and Atmospheric Administration climate database.

### Molecular Analysis

For the swab DNA extractions, we used the PrepMan® Ultra Sample Preparation Reagent (Applied Biosystems, Foster City, CA, USA) that is designed to rapidly extract fungal and bacterial DNA from complex samples (see Supplementary information for full protocol) ^15,69,74^. We extracted genomic DNA from all skin biopsies using the GeneJET Genomic DNA Purification Kit (K0722; Thermo Scientific). To detect *Oo* via qPCR, we used the primers and probe designed by Allender et al. ^75^. Triplicate reactions were run in 25 μL total volumes, which were composed of 12.5 μL 2X TaqMan Fast Advanced Master Mix (Applied Biosystems), 2.75 μL nuclease-free water, 0.9 μM of each primer, 0.25 μM of probe, 400 ng BSA, and 5 μL of non-diluted extracted DNA (variable concentrations). For our standards, we used a dynamic range of 1 x 10^−1^ to 1 x 10^−5^ ng/ μL of gDNA and substituted gDNA with nuclease-free water for the negative controls. We used the following fast cycling parameters: initial incubation at 95°C for 20 s, followed by 50 cycles of 95°C for 1 s, and 60°C for 20 s (data capture step) on a QuantStudio 3 0.1 μL block real-time instrument (Applied Biosystems).

Previous studies have used internal transcribed spacer 1 (ITS1) copy number estimates ^75^ and a dynamic range of known DNA concentrations ^76^ to infer positive *Oo* infection and copy number via qPCR. The lower limit of sensitivity for the assay proposed by Allender et al. ^75^ was ~10 ITS1 copies. However, ITS1 copy number variation among strains and isolates has been documented in other fungal pathogens ^77^, and currently, there are no published data indicating whether or not *Oo* ITS1 copy number varies across strains or regional isolates of *Oo*. The assay proposed by Bohuski et al. ^76^ utilized a 10-fold serial dilution of genomic DNA from 500,000 fg to 50 fg, and their reported detection limit was 5 fg. The latter approach, however, does not provide a quantity for the number of nuclear equivalents in solution.

Since the development of these initial assays, the *Oo* genome has been sequenced and the haploid genome size is estimated to be 21.9 Mb ^78^. Therefore, we combined the methods utilized by Allender et al. ^75^ and Bohuski et al. ^76^ by using genome size as an estimate of the number of nuclei in solution (i.e., genomic equivalents). To estimate genomic equivalents, we updated the equation that Allender et al. ^75^ employed by substituting the molecular mass of the ITS1 region with the molecular mass of the nuclear genome. Therefore, we estimated the mass of one *Oo* genomic equivalent at approximately 2.36 x 10^−5^ ng or 23.6 fg using the following equation:

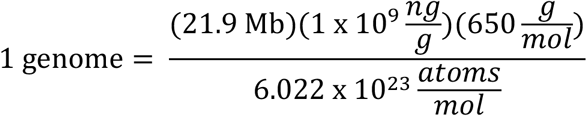

Our positive controls consisted of 10-fold serial dilutions spanning a dynamic range of 500,000 fg to 50 fg (i.e., ~21,186 genomic equivalents to ~2.1186 genomic equivalents). We conducted reactions using duplicate positive and negative controls. To compensate for increased variance at the lower concentrations of the dynamic range, the 500 fg and the 50 fg standard reactions were run in quadruplicate.

### Statistical Analysis

Using the equation: mass/SVL^2^, we calculated body condition index (BCI) ^18,79,80^. We also estimated prevalence among counties, species, and across our total sample size. We estimated 95% binomial confidence intervals with a logistic parameterization for all categorical data (i.e., county and species) using the R package “binom” ^81^. We compared the skin biopsy qPCR *Oo* detections to the swab detections and estimated the false-negative rate from swabs as well as the false negative-rate of skin biopsies using the R package “cutpointr” ^82^. We then used Bayesian estimates of “corrected” *Oo* prevalence with 95% credible intervals and used our false-negative rate estimate to set our informative priors for lower sensitivity rate ^83^. We conducted four Fisher’s Exact Tests to examine the associations between: 1) SOI and *Oo* occurrence; 2) species and SOI; 3) sex and *Oo* occurrence; and 4) life stage and *Oo* occurrence. We were concerned with comparison-wise error only; therefore, we did not apply correction to *P*-values obtained from our analyses ^84^. We also used a Kruskal-Wallis non-parametric analysis to test for differences in median BCI values of snakes with positive and negative qPCR results as well as differences in BCI and SOI. To minimize the effects of snake size disparity between life stages and across species, we tested for differences in BCI values only within each life stage, and within species, but not across life stages or species.

To evaluate the contributions of predictive factors to the probability of the *Oo* occurrence, we constructed 14 logistic regression mixed effects models (two intercept-only models) with the binomial family distribution using the “glmmTMB” package ^85^. We considered the following explanatory variables: species, life stage, mean annual temperature, and mean annual precipitation. Eight of the models included sample site as a random effect, as well as sample month (included as a first order autoregressive [AR1] variable), thus accounting for temporal autocorrelation in the model. Six models included only sample month as an AR1 variable. To avoid overparameterization, we constructed the models with only one or two fixed effects. We then computed AICc for each candidate model to determine the most parsimonious model using the “MuMIn” package ^86^. Our analyses did not consider the effects of variable interactions owing to low sample size. We were unable to collect snakes during several months of the 2019 summer season and most of the 2020 spring season, and thus, we did not estimate seasonal *Oo* prevalence. Using the most parsimonious AICc model, we inferred infection probability via estimated marginal means (EMM) obtained from an inverse logit transformation using the “emmeans” package ^87^. *Post hoc* analysis of the EMM for each predictor of was conducted using Tukey’s HSD method.

All statistical analyses were conducted using R (R version 3.6.1, www.r-project.org, accessed 5 July 2019). We used the Thermo Fisher Connect™ Cloud Dashboard Software (Thermo Fisher Scientific) for qPCR data processing and analysis, and we created distribution maps using QGIS open-source software (QGIS version 3.16.3, QGIS Development Team, QGIS Geographic Information System, Open Source Geospatial Foundation, https://www.qgis.org/en/site/, accessed 15 Jan 2019).

### Data availability

The raw data generated and analyzed for this study are available from the Dryad depository (https://doi.org/XX.XXXX/dryad.XXXXXXX).

## Acknowledgments

We thank Dustin McBride for logistical advice. We thank Jeremiah Leach, Daniel Puckett, Gabriella Solis, Thomas Marshall, Shashwat Sirsi, Michaela Bowlsby, Cheyenne N. Gonzales, Arya J. Sanjar, Juliette Garza, Robert Tyler, Stephen Roussos, Jeremy Weaver, Mark Pyle, Kasey Jobe, and Nathan Rains for assistance the field. We thank Toriann Molis, Mario B. Sarmiento, Clarissa Rivera, Gabriella Solis, Mireya A. Escandon, Stephanie Monroe, Carlos Baca, and Rebecca M. Brunner for assistance in the laboratory. This project was funded by Texas Parks and Wildlife Department and the U.S. Fish and Wildlife Service through the State Wildlife Grant Program (TX T-164-R-1 to D.R. and M.R.J.F.), the Conservation License Plate Program (contract #531961 to S.J.M.), and startup funds to D.R. from Texas State University.

## Author Contributions

D.R., P.C., and M.R.J.F. conceived and designed the study. S.F.H., S.J.M., M.R.J.F. and, D.R. attained funding for the study. S.F.H., J.Y., P.C., S.J.M., M.R.J.F., and D.R. participated in surveys and sample collection. S.F.H. and D.R. performed molecular diagnostics. S.F.H., C.G.B., and, P.C. designed and performed the statistical analyses. S.F.H. and D.R. co-wrote the draft and constructed the figures. S.F.H., C.G.B., J.Y., P.C., M.R.J.F., S.J.M., and, D.R. contributed to the interpretation of the results and writing the final version of the manuscript.

