## Supplementary for "Comparative host-pathogen dynamics of Snake Fungal Disease in sympatric species of water snakes (*Nerodia*)"

**Tables**

**Table S1.** Summarized results from AIC model selection conducted on 14 models

**Table S2.** Summary of model selection results for the four best fit models

**Table S3.** Summarized output of our most parsimonious model

**Table S4.** List of predicted probability of infection for species

**Table S5.** List of predicted probability of infection for life stage

**Table S6.** List of predicted probability of infection for species and life stage

**Table S7.** Summarized criteria for life stage classification of three *Nerodia* sp.

**Figures**

**Figure S1.** Map showing relative sample size and per site species composition

**Figure S2.** Binned residual plot generated from most parsimonious model

**Figure S3.** Quantile-quantile plot for most parsimonious model

**Methods**

DNA extraction from swabs

Logistic regression model diagnostics

**Results**

Logistic regression model diagnostics

**Appendix S1. List** of sampled counties, sites within counties, qPCR results for each site, and swab sample size for each size.

### Tables

**Table S1.** Summary statistics from tests of 14 models constructed to estimate variable effects on *Ophidiomyces ophiodiicola* detection (y), their AIC value, and the number of parameters for each model (K), delta values (Δ), and model weights (w). The random effects are denoted in parentheses. First order autoregression applied to random effects are indicated by "ar1". Site = sampled site; Month = the month when sampling event occurred; Year = calendar year for sampling event; LS = snake life stage; Species = snake species; SVL = snout-vent-length measured in centimeters; Temp = mean monthly low temperature for each sample site; Prec = mean monthly precipitation for each site.

| Model | AICc | K | Δ | *w* |
|---|---|---|---|---|
| y = Species + LS + ar1(Month + 0 \| Year) | 176.12 | 7 | 0.00 | 0.36 |
| y = LS + ar1(Month + 0 \| Year) | 176.59 | 5 | 0.47 | 0.28 |
| y = Species + LS + (1 \| Site) + ar1(Month + 0 \| Year) | 177.25 | 8 | 1.13 | 0.20 |
| y = LS + (1 \| Site) + ar1(Month + 0 \| Year) | 177.94 | 6 | 1.82 | 0.14 |
| y = Species + SVL + ar1(Month + 0 \| Year) | 183.73 | 6 | 7.61 | 0.01 |
| y = Species + SVL + (1 \| Site) + ar1(Month + 0 \| Year) | 184.37 | 7 | 8.25 | 0.01 |
| y = 1 + (1 \| Site) + ar1(Month + 0 \| Year) | 200.07 | 4 | 23.95 | 0.00 |
| y = 1 + ar1(Month + 0 \| Year) | 200.49 | 3 | 24.37 | 0.00 |
| y = Species + (1 \| Site) + ar1(Month + 0 \| Year) | 200.62 | 6 | 24.50 | 0.00 |
| y = Species + ar1(Month + 0 \| Year) | 200.95 | 5 | 24.83 | 0.00 |
| y = Temp + (1 \| Site) + ar1(Month + 0 \| Year) | 202.15 | 5 | 26.03 | 0.00 |
| y = Prec + (1 \| Site) + ar1(Month + 0 \| Year) | 202.19 | 5 | 26.07 | 0.00 |
| y = Temp + Prec + (1 \| Site) + ar1(Month + 0 \| Year) | 202.55 | 6 | 26.43 | 0.00 |
| y = Temp + Prec + ar1(Month + 0 \| Year) | 204.30 | 5 | 28.18 | 0.00 |

**Table S2.** Summary of model selection results for the four best fit models. The number of parameters (K), log-likelihood (*LL*), AIC score corrected for small sample size (AICc), delta (Δ), weight (*w*) and parameter coefficients with standard errors are listed for each model. Parameter estimates are relative to adult *Nerodia harteri harteri*—or all adults in the model that did not include species. LS = snake life stage.

| | | | | | | | β | | | |
|---|---|---|---|---|---|---|---|---|---|---|
| **Model** | **K** | ***LL*** | **AICc** | **Δ** | ***W*** | **Intercept** | ***N. e. transversa*** | ***N. rhombifer*** | **juvenile** | **sub-adult** |
| Species + LS + ar1(Month + 0 \| Year) | 7 | -80.67 | 176.12 | 0 | 0.36 | 1.56 ±0.67* | -1.00 ±0.66 | -1.28 ±0.64* | -2.87 ±0.66*** | -2.00 ±0.57*** |
| LS + ar1(Month + 0 \| year) | 5 | -83.09 | 176.59 | 0.47 | 0.28 | 0.62 ±0.37 | - | - | -2.69 ±0.60*** | -1.71 ±0.50*** |
| Species + LS + (1 \| Site) + ar1(Month + 0 \| Year) | 8 | -80.11 | 177.25 | 1.13 | 0.20 | 1.59 ±0.77* | -1.10 ±0.71 | -1.45 ±0.71* | -2.85 ±0.67*** | -2.01 ±0.59*** |
| LS + (1 \| Site) + ar1(Month + 0 \| Year) | 6 | -82.68 | 177.94 | 1.82 | 0.14 | 0.49 ±0.44 | - | - | -2.65 ±0.62*** | -1.71 ±0.52*** |

* estimates with a P-value < 0.05

**Table S3.** Summarized conditional model output from our most parsimonious model showing parameter coefficients, standard error (SE), z-score (z), and *P*-value (p). All parameter estimates are relative to *Nerodia harteri* adults.

| | Estimate | SE | Z | p |
|---|---|---|---|---|
| Intercept | 1.562 | 0.675 | 2.313 | 0.021 |
| Species: *erythrogaster* | -0.998 | 0.658 | -1.516 | 0.130 |
| Species: *rhombifer* | -1.280 | 0.639 | -2.003 | 0.045 |
| Stage: juvenile | -2.874 | 0.655 | -4.387 | 0.00001 |
| Stage: sub-adult | -2.002 | 0.570 | -3.515 | 0.0004 |

*Time series: Variance=0.798, Corr(ar1)=0.72, N=150.*

**Table S4.** Summary of the predicted probabilities of *Ophidiomyces ophiodiicola* infection for each species associated with the standard error (SE), degrees of freedom (df), lower confidence limit (L), upper confidence limit (U), t-ratio (t), and *P*-value (p). Probabilities are averaged over the factor: life stage.

| Species | prob | SE | df | L | U | t | p |
|---|---|---|---|---|---|---|---|
| harteri | 0.484 | 0.149 | 143 | 0.259 | 0.716 | -0.106 | 0.916 |
| erythrogaster | 0.257 | 0.117 | 143 | 0.111 | 0.489 | -1.730 | 0.085 |
| rhombifer | 0.207 | 0.094 | 143 | 0.092 | 0.403 | -2.339 | 0.021 |

**Table S5.** Summary of the predicted probabilities of *Ophidiomyces ophiodiicola* infection for each life stage. The standard error (SE), degrees of freedom (df), lower confidence limit (L), upper confidence limit (U), t-ratio (t), and *P*-value (p). Probabilities are averaged over the factor species.

| stage | prob | SE | df | L | U | T | p |
|---|---|---|---|---|---|---|---|
| juvenile | 0.112 | 0.067 | 143 | 0.040 | 0.277 | -3.081 | 0.003 |
| subadult | 0.232 | 0.113 | 143 | 0.096 | 0.462 | -1.898 | 0.059 |
| adult | 0.691 | 0.109 | 143 | 0.489 | 0.839 | 1.571 | 0.1183 |

**Table S6.** Summary of the predicted probabilities of *Ophidiomyces ophiodiicola* infection for each species at each life stage, standard error (SE), degrees of freedom (df), lower confidence limit (L), upper confidence limit (U), t-ratio (t), and *P*-value (p).

| Species | stage | prob | SE | df | L | U | t | p |
|---|---|---|---|---|---|---|---|---|
| harteri | adult | 0.827 | 0.097 | 143 | 0.609 | 0.936 | 2.313 | 0.022 |
| erythrogaster | adult | 0.637 | 0.150 | 143 | 0.375 | 0.837 | 0.868 | 0.387 |
| rhombifer | adult | 0.570 | 0.132 | 143 | 0.352 | 0.764 | 0.524 | 0.601 |
| harteri | juvenile | 0.212 | 0.126 | 143 | 0.072 | 0.484 | -1.743 | 0.084 |
| erythrogaster | juvenile | 0.090 | 0.065 | 143 | 0.026 | 0.267 | -2.941 | 0.003 |
| rhombifer | juvenile | 0.070 | 0.048 | 143 | 0.021 | 0.205 | -3.473 | 0.001 |
| harteri | subadult | 0.392 | 0.165 | 143 | 0.170 | 0.670 | -0.635 | 0.527 |
| erythrogaster | subadult | 0.192 | 0.112 | 143 | 0.067 | 0.441 | -1.983 | 0.049 |
| rhombifer | subadult | 0.152 | 0.098 | 143 | 0.049 | 0.386 | -2.265 | 0.025 |

**Table S7.** Summarized snout-to-vent length measurements used to determine life stage for *Nerodia rhombifer*, *Nerodia erythrogaster transversa*, and *Nerodia harteri harteri* collected in the Brazos River Drainage, Texas. All measurements are in centimeters (SM = sexual maturity; M = male; F = female).

| Species | Juvenile | Sub-Adult | Adult M | Adult F | Source |
|---|---|---|---|---|---|
| *N. rhombifer* | < 30.0 | 30.0 - SM | ≥ 47.0 | ≥ 67.0 | Kofron 1979, Plummer 1992; Aldridge et al. 1995 |
| *N. e. transversa* | < 30.0 | 30.0 - SM | ≥ 55.0 | ≥ 66.5 | Gibbons and Dorcas 2004; Lacki et al. 2005 |
| *N. h. harteri* | < 30.0 | 31.0 - SM | ≥ 38.0 | ≥ 46.0 | Green et al. 1994; Greene et al. 1999 |

#### Figures

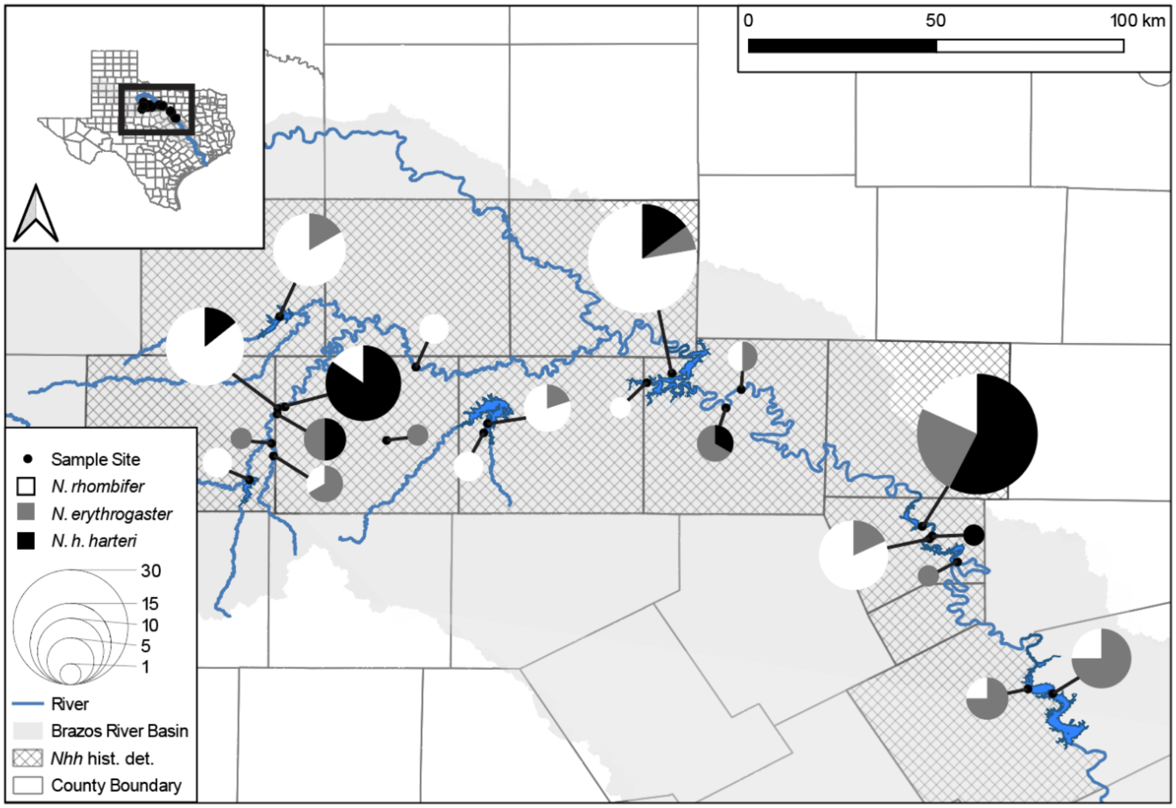


**Figure S1.** *Nerodia* captures in the upper Brazos River basin. The size of the pie chart indicates the relative sample size of snakes captured at each site. Counties shaded with cross hatching are counties represent the historical range of *Nerodia harteri harteri*.

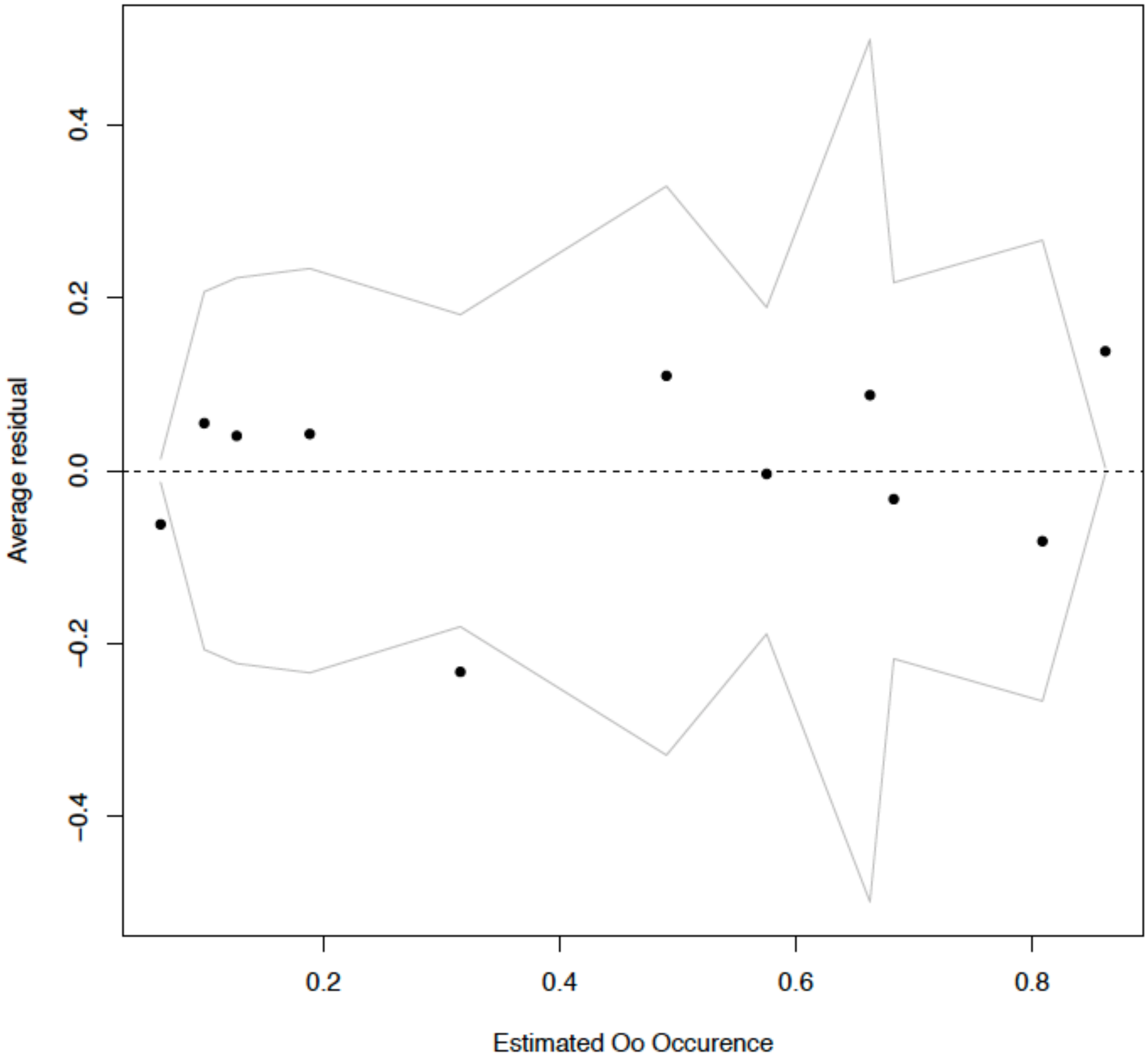


**Figure S2.** Binned average residuals with 95% confidence interval of logistic regression mixed effects model for the effect of three *Nerodia* species and three life stages on *Ophidiomyces ophiodiicola* occurrence.

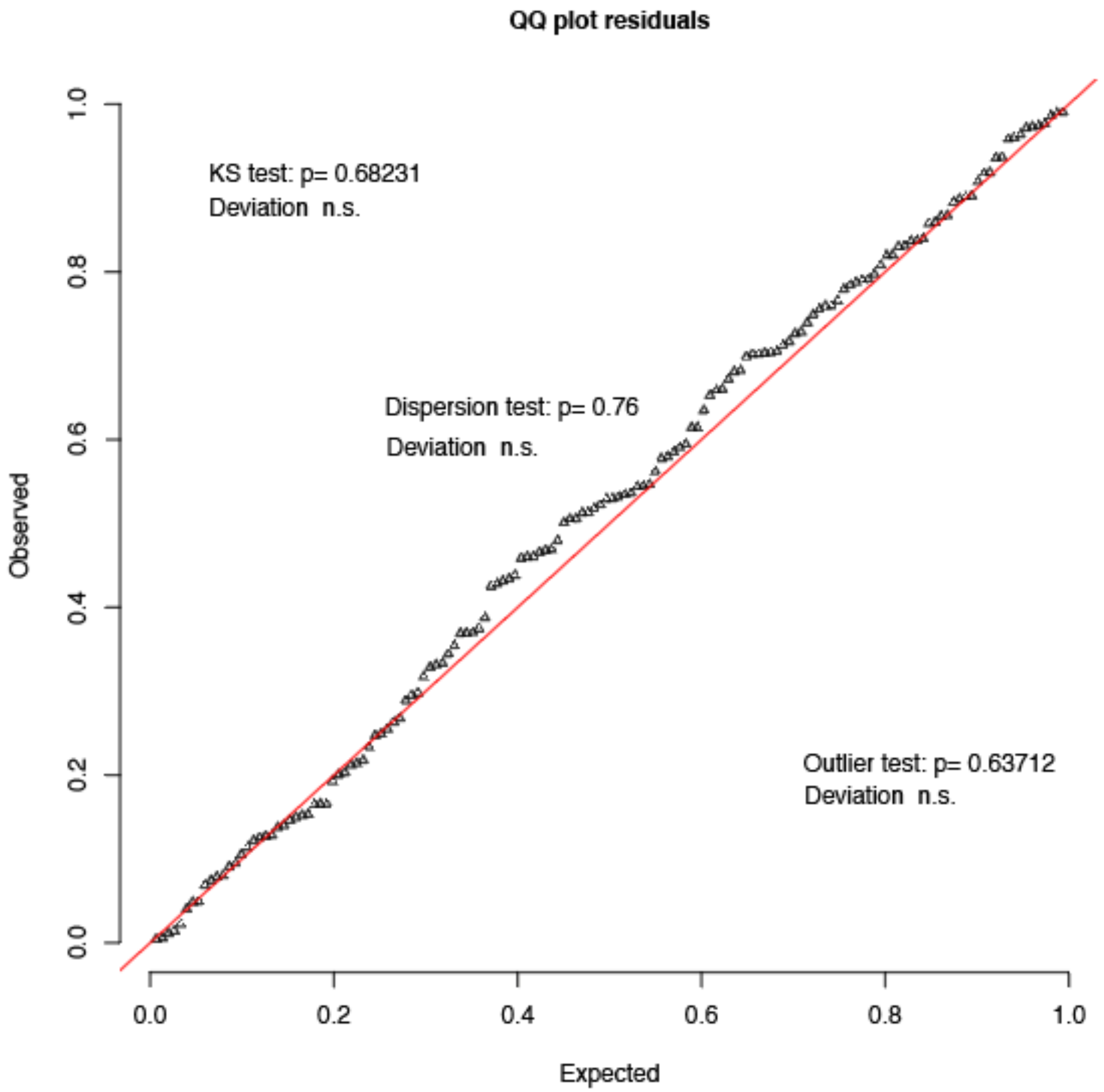


**Figure S3.** Quantile – quantile residual plot constructed from logistic regression mixed effects model for the effect of three *Nerodia* species and three life stages on *Ophidiomyces ophiodiicola* occurrence. Results (p-value) from a KS test for non-parametric distribution, dispersion test, and outliers are overlaid on plot (n.s. = not significant).

### Methods

#### DNA Extraction Protocol

We added 50 µL of PrepMan™ Ultra (Applied Biosystems) to each 2 mL screw cap centrifuge tube containing the swab sample. We placed the tubes into a floating foam tube rack and immersed the tubes in a water bath at 100 ºC for 10 mins. After, we allowed the samples to cool for 2 mins. Then we centrifuged the samples at 12,000 RCF for one minute. Using sterilized forceps, we inverted the swab inside the tube to orient the applicator towards the screwcap. To separate the supernatant from the applicator, we centrifuged the samples at 12,000 RCF for one minute. After which, we discarded the swab. We then centrifuged the supernatant at 12,000 RCF for 10 minutes to pellet precipitates and debris in solution. Lastly, we recovered ~30 µL of the supernatant and placed into a labeled, sterile 1.5 µL microcentrifuge tube. Extracted DNA was stored at -20 ºC prior to qPCR analyses.

#### Logistic Regression diagnostics and goodness of fit

To diagnose goodness of fit for our mixed effects model, we calculated the likelihood-ratio based pseudo-$R^2$ value and the likelihood-ratio based pseudo-$R^2$ value with a Nagelkerke modification (Nagelkerke 1991) using the "MuMIn" package (Barton 2019). We also calculated the Pearson $X^2$ and log-likelihood approximated deviance goodness of fit statistics (Forthofer et al. 2007). To assess deviations from the expected distribution, we conducted a Kolmogorov-Smirnov test, a dispersion test, and outlier test using the package "DHARMa" (Hartig 2020). We visually inspected the distribution of the residuals by constructing a plot of the binned residuals (Gelman and Hill 2006). Lastly, we constructed a quantile-quantile plot using the package "DHARMa". All analyses were conducted using R software (R Core Team 2019).

### Results

#### Logistic Regression diagnostics and goodness of fit

The pseudo-$R^2$ and modified pseudo-$R^2$ values were 0.197 and 0.272, respectively. The Pearson goodness of fit and the deviance statistics were 0.752 and 0.140, respectively. The distribution of the binned residuals contained 73.0% of the observations within the ± 2 SE bands (Figure S2). The p-values from the KS, dispersion, and outlier tests were not significant (p = 0.682, 0.760, and 0.637; Figure S3). The quantile-quantile plot did not exhibit any significant outliers (Figure S3).

**Appendix S1.** List of counties sampled for *Nerodia*, the sample site within the county, the number of positive *Oo* qPCR tests, the number of negative qPCR tests, and the total number of swabs tested for each site (n).

| County | Site | Positive | Negative | n |
|---|---|---|---|---|
| Bosque | Morgan Lake Side Park | 3 | 1 | 4 |
| Haskell | Lake Stamford | 8 | 4 | 12 |
| Hill | Lake Whitney State Park | 2 | 6 | 8 |
| Hood | City Park | 3 | 8 | 11 |
| | DeCordova Bend Park | 0 | 1 | 1 |
| | Hunter Park | 20 | 13 | 33 |
| | Private Property | 1 | 0 | 1 |
| Jones | Doby Ranch | 2 | 1 | 3 |
| | Fort Phantom Hill | 1 | 1 | 2 |
| | Threemile Creek | 0 | 1 | 1 |
| Palo Pinto | Eagle Creek | 0 | 3 | 3 |
| | Possum Kingdom State Park | 0 | 1 | 1 |
| | Rochelles Canoe Rental | 1 | 1 | 2 |
| | Graham | 13 | 14 | 27 |
| Shackelford | Lueders | 11 | 3 | 14 |
| | Creek Park | 0 | 1 | 1 |
| | Fort Griffin State Park | 0 | 2 | 2 |
| | Hog Creek | 1 | 12 | 13 |
| | Post Oak Creek | 0 | 4 | 4 |
| Stephens | Bob Clark Landing | 3 | 2 | 5 |
| | Private Property | 0 | 2 | 2 |
| | | 69 [a] | 81 [a] | 150 [a] |

[a] Total
